# Electrocortical Evidence for Long-Term Incidental Spatial Learning Through Modified Navigation Instructions

**DOI:** 10.1101/280842

**Authors:** Anna Wunderlich, Klaus Gramann

## Abstract

The use of Navigation Assistance Systems for spatial orienting has become increasingly popular. Such automated navigation support, however, comes with a reduced processing of the surrounding environment and often with a decline of spatial orienting ability. To prevent such deskilling and to support spatial learning, the present study investigated incidental spatial learning by comparing standard navigation instructions with two modified navigation instruction conditions. The first modified instruction condition highlighted landmarks and provided additional redundant information regarding the landmark (contrast condition), while the second highlighted landmarks and included personal relevant information regarding the landmark (personal-reference condition). Participants’ spatial knowledge of the previously unknown virtual city was tested three weeks later. Behavioral and electroencephalographic (EEG) data demonstrated enhanced memory performance for participants in the modified navigation instruction conditions without further differentiating between modified instructions. Recognition performance of landmarks was better and the late positive complex of the event-related potential (ERP) revealed amplitude differences reflecting an increased amount of recollected information for modified navigation instructions. The results indicate a significant long-term spatial learning effect when landmarks are highlighted during navigation instructions.

## Introduction

Successful orienting in and navigating through the environment is one of the most important abilities of living creatures facing daily challenges in dynamic environments. Being an essential part of nearly all indoor and outdoor activities, spatial orientation is closely connected to general memory processes. However, technological developments in the last decades have changed the demands on individual navigation abilities and the way we memorize space. The use of motorized vehicles increased the speed of movement and, consequently, the spatial scale of the navigated environment. Using motorized vehicles also requires control processes that allow save driving and interaction with other road users while keeping with traffic regulations. Navigation assistance systems were developed to help the driver in this multi-tasking situation and to provide the necessary directions enabling the driver to focus on the driving task while navigating to a goal location.

Due to these benefits, navigation assistance systems based on Global Positioning (GPS) are now widely used even when walking through familiar areas. With this growing trust in automation, new issues can be observed that are typically associated with automated systems. One important issue is over-trust [1, 2]: If a person is responsible for multiple tasks, and in addition, has to monitor an automated task that is based on seemingly reliable information, s/he tends to trust too much in the automation. In other words, in an orientation context the user outsources the navigation task completely to the assistance system [3] leading to decreased processing of environmental information including landmarks (salient orienting points) and their spatial relations. As a consequence, spatial knowledge acquisition degrades and spatial orienting skills decline with the use of navigation assistance systems [4].

Different methods have been tested to overcome the negative impact of navigation assistance. Such approaches aim at enhancing spatial learning by modifying the kind of spatial information and/or the modalities (tactile, visual and/or auditory) of navigation instructions provided to the user (see [5] and [6] for an overview). Using the central role of landmarks in spatial orienting, Gramann and colleagues [5] demonstrated improved spatial learning with modified navigation instructions. In their study, participants were assigned to one of three conditions that differed in the amount of information provided with the auditory navigation instruction. Compared to a control condition (“Please turn left at the junction.”), the two modified navigation instruction conditions highlighted landmarks at navigation-relevant intersections (right or left turn) and provided additional information related to this landmark. This was either a short description of the landmark (e.g., “Please turn left in front of the embassy. Here, an ambassador works.”) or personally relevant information (e.g., “Please turn left in front of the embassy. This is the embassy of your favorite country ‘New Zealand’.”). The acquired spatial knowledge was evaluated directly after the navigation phase demonstrating improved spatial knowledge in both modified navigation instruction conditions as compared to the control group while the mental load for all instruction conditions was the same.

Spatial knowledge acquisition can be described at different levels. According to the model introduced by Siegel and White [7], a first level describes Landmark Knowledge representing knowledge about objects in space. A sequential connection between those landmarks in space is described by Route Knowledge. A complex, map-like Survey Knowledge then includes spatial relations between landmarks enabling complex online and offline computations including short-cuts using previously unknown routes. While the original model proposes a sequential development from landmark to survey knowledge, several authors argue that the acquisition of spatial knowledge on different representational levels takes place in parallel [9–11]. In all accounts, however, encoding of landmarks and subsequent integration of the same into a spatial representation is essential to spatial knowledge acquisition. To experimentally address the encoding stage during spatial knowledge acquisition, the Levels of Processing (LoP) approach [11] offers a theoretical framework. Because navigation does not primarily demand semantic processing of the perceptible environment (e.g., different buildings at a navigation-relevant junction), a cue augmenting an otherwise unnoticed building at a navigation-relevant intersection, attracts attention towards this building and renders a previously meaningless object salient and relevant. As a consequence, augmenting buildings by navigation instructions can initiate a deeper elaboration of the presented information that is associated with a spatial location. Following the LoP model, this elaboration of specific landmark information should be associated with a longer maintenance of all information connected to the landmark and the surrounding. Moreover, the self-reference effect (for a review see [12]) expands the elaboration of landmark information further and increases the depth of processing strengthening the memory trace [13]. Using navigation instructions referring to landmarks might thus lead to incidental spatial learning [14].

Based on the significant short-term improvement in spatial knowledge acquisition in the previous study [5], the current study used behavioral and neural measures to investigate whether modified navigation instructions can also lead to long-term improvement of spatial knowledge. Beyond performance in navigation-related tasks, brain electrical activity recorded with EEG was used to investigate components of the event-related potential (ERP). The analysis focused on the late positive complex (LPC) of the ERP which is assumed to reflect information recollection (for a review see [15]). The amplitude of the posterior LPC increases with an increasing amount of information that is recollected [16, 17]. Based on the findings of Gramann and colleagues [5], we expected improved performance even after three weeks when using modified navigation instructions to navigate an unknown environment. In addition, increased LPC-amplitudes were expected for augmented landmarks with the most pronounced amplitudes for the personal-reference navigation instruction condition.

## 2 Materials and Methods

### 2.1 Participants

Forty-two participants with at least two years ownership of a driver license participated in the study. Participants were recruited through an existing database or direct contact. All volunteers received 10 Euro/h allowance for their participation. Gender distribution was kept constant for all conditions. For the analyses, 21 female and 21 male participants were included with an average age of 26.62 years (*SD* = 4.36 years, *Min* = 19 years, *Max* = 37 years). All had normal or corrected to normal vision and gave informed consent prior to the study. The study was approved by the local research ethics committee.

### 2.2 Measurements and Apparatus

The experiment took place in a driving simulator at TU Berlin consisting of a stripped VW Touran driver’s seat environment and a projector screening the scene onto a white wall approximately two meters in front of the driver. The projection had a horizontal and vertical viewing angle of 90 degrees and 54 degrees, respectively. These angles depended on the individual body size (see figure 1A). The steering and pedal behavior was recorded through linking the seat box to the Game Controller, MOMO Racing Force Feedback Wheel (Logitech, Switzerland). This integration limited a realistic driving experience to a certain degree by reducing the maximum angle of the steering wheel to 110 degrees. To compensate for this, the sensitivity of the control was increased. The Game Controller further provided two pedals to simulate an automatic transmission.

**Figure 1:**
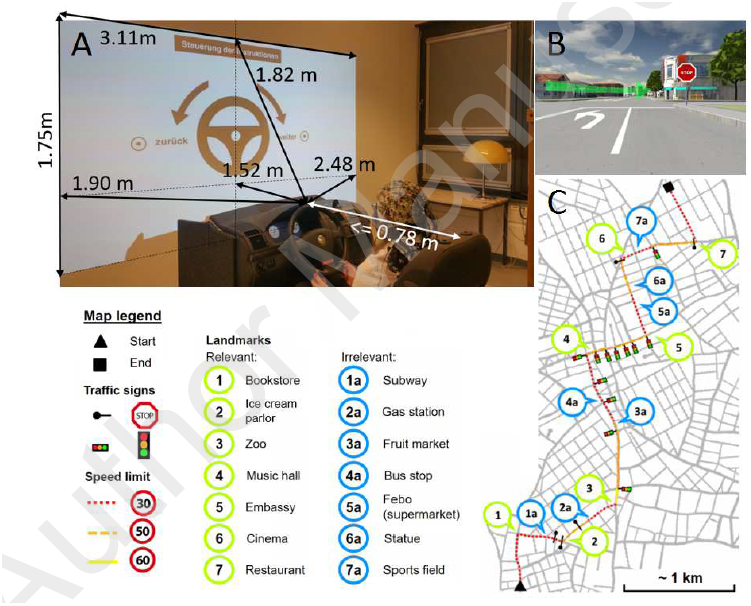
**A)** Setup of the experiment with driving simulator and projection area and the important distance measurements **B)** The visual navigation instruction is a semitransparent hologram arrow projected in the environment at decision points. **C)** Map of the virtual city with the route and marker for the relevant and irrelevant landmarks and traffic details.

The virtual city model from Gramann et al. [5] was used, based on the open source simulator software OpenDS (http://www.opends.de). The size of the city was approximately 36km^2^ and the simulation restricted traffic to one single car at the beginning of the experiment to avoid an impact of traffic on the processing of environmental information. The route headed from one suburban quarter through an inner-city area to another suburban quarter, where the route ended. Along the way, different speed limits and stop signs or traffic lights were present and the route included seven direction changes at intersections with an unpredictable order of turning direction. Salient buildings that contrasted with the surrounding buildings, hereafter named landmarks, were placed in the virtual environment. Dependent on their location along the route, landmarks were categorized as either relevant (intersection with direction change) or irrelevant (straight route segment) for successful navigating (see figure 1C). The number of irrelevant landmarks between two direction changes varied from 0 to 2 (*M* = 0.88; *SD =* 0.78). The location of relevant landmarks at the intersections provided no indicator for turning directions and navigation instructions were triggered whenever the vehicle passed a predefined point 100 meters ahead of a direction change. Navigation instructions consisted of an auditory instruction (e.g., “In 100 meters, turn right at the next intersection”) and, in addition, a visually presented arrow that simulated a head-up display (a semitransparent hologram arrow projected onto the environment; see Figure 1B). The augmented route guiding system with auditory instructions allowed participants to continuously direct their attention towards the environment [18, 19].

In case the participant left the predefined route, an automated resetting mechanism stopped the car was positioned back on the correct route facing the last intersection.

### 2.3 Study Design and Procedure

The experiment had a 3×2 factorial mixed measures design, with three different navigation instruction conditions (standard, contrast modified, and personal-reference modified) as between subject factor and a repeated measures factor experimental session (first and second navigation session) in which participants first navigated the route according to the navigation instructions and in the second session had to solve a series of spatial tasks as described below. The two sessions were separated by a three-week period (*M* = 21.07 days, *SD* = 1.65 days, *Min* = 20 days, *Max* = 24 days) to investigate the long-term impact of modified navigation instructions on spatial learning and memory. Participants were not aware that they would be tested on the same environment in the second session.

#### Pretest Questionnaire

Every participant, irrespective of the subsequent navigation instruction condition, had to answer a pretest questionnaire about her personal preferences regarding music, literature and other dimensions via e-mail. The answers were included in the auditory navigation instructions via text to speech software (Voice RSS, http://www.voicerss.org) for all participants, who were randomly assigned to the personal-reference condition.

Within the test sessions, participants answered several questionnaires between the tasks on a tablet (iPad 1, Apple Inc., Curtino, California using the software LimeSurvey, Hamburg, Germany). Overall, the first experimental session lasted approximately one hour and included the tasks described in the following.

#### Navigation Drive

Participants were instructed to abide traffic rules and were given 5 minutes to drive in a different environment to get accustomed to the wheel and pedal setup. Afterwards they were placed in the test environment and asked to follow the navigation instructions during driving. Participants’ mental load during the navigation-guided drive was assessed using subjective (unweighted National Aeronautics and Space Administration Task Load Index, NASA-TLX, [20]) and objective measurements (driving parameters including velocity, distance to ideal line, pedal and steering wheel changes). Afterwards, the affective state of the participant was assessed by the Affect Grid [21] and three questions on a four-point Likert-scale regarding simulation sickness (nausea, headache, and dizziness).

#### Assessment of Individual Spatial Abilities

The first session included also several questionnaires focusing on driving and navigation habits, gaming experience, and spatial orientation (Santa Barbara Sense of Direction, SBSOD, [22]) as well as the Reference Frame Proclivity Test (RFPT) assessing the individual preference to use an egocentric or an allocentric reference frame during a path integration task [23, 24].

The second experimental session aimed at quantifying participants’ spatial knowledge about the virtual environment that had been experienced during the first navigation session three weeks prior. Participants worked on five tasks measuring different levels of spatial knowledge in the following order.

#### Landmark Recognition Task

The landmark recognition task was designed to allow the analysis of event-related potentials and presented participants with snap-shots of 21 intersections from the test environment in the drivers’ perspective in a random order. For every shown junction, participants had to decide whether the landmark was relevant, or irrelevant, or had not been present (novel landmark). For relevant landmarks, participants were instructed to steer into the direction that was instructed during the first navigation session. Furthermore, participants were instructed to push the gas pedal for irrelevant landmarks and to apply the brakes in case of novel landmarks. Overall, seven landmarks for each of the three landmark types (relevant, irrelevant, novel) were presented. The task included ten repetitions of all landmark stimuli to get a sufficient number of epochs for the analysis of event-related potentials. The landmark recognition task focused on landmark knowledge. However, because the responses required navigation decisions it also tested route knowledge (or Heading Orientation according to [25]) including stimulus-response associations. We expected improved landmark and route recognition for the modified navigation instruction conditions.

#### Scene Sorting Task

This task did also address landmark and route knowledge but was more focusing on the route representation as sequence of landmarks. Participants were asked to select the relevant and irrelevant landmarks out of all 21 landmarks (the same as in the landmark recognition task) and place them in the correct chronological order encountered during the first navigation session.

#### Sketch Mapping 1

Afterwards, participants were instructed to draw a map of the virtual environment including the route driven as well as any buildings, traffic lights, signs, or any other route-connected information the participant remembered on an empty DIN A3 page. This task combined all levels of spatial knowledge with a specific focus on survey knowledge.

#### Navigation Drive without Assistance

Based on the assumption that the recall of information is best when learning and test situation are congruent [26], this task setting was identical to the learning situation. Participants were instructed to drive the identical route as in the first navigation session without navigation assistance. Participants got accustomed with the simulator setup before starting the task in a five-minute training session. Afterwards, the car was placed at the start of the route. In case of incorrect turns, the car was automatically set back to the last junction before the incorrect direction change. After reaching the destination, the NASA-TLX, Affect Grid, and simulation sickness questionnaire were filled out.

#### Sketch Mapping 2

Subsequently, participants were asked to draw a sketch map of the virtual environment. The objective of this second map was to reveal differences in recovering survey knowledge after being confronted with the environment for a second time.

In a final questionnaire, participants provided feedback about the experiment and the modified navigation instructions.

### 2.4 Electroencephalography (EEG)

The EEG was recorded continuously for the complete second session using 64 channels (BrainAmps, Brain Products, Gilching, Germany) positioned in an elastic cap (EASYCAP, Herrsching, Germany) according to the extended 10% system [27]. All electrodes were referenced to FCz and the data was collected with a sampling rate of 500 Hz, band-passed from 0.016 Hz to 250 Hz. One electrode below the left eye (vEOG) was used to record vertical eye-movement. Time synchronization and disk recording of the data streams from the driving simulator and EEG was done using Lab Streaming Layer (LSL, [28]). Age of sample was corrected in the analyses by subtracting 50ms for the delay induced by the EEG amplifier setup^1^ and another 30ms for the input lag of the projector^2^.

For data processing, the interactive Matlab toolbox EEGLAB [29] was used. The raw data was high pass filtered at 0.5 Hz, low pass filtered at 75 Hz, and down sampled to 250 Hz. Subsequently, noisy channels and artifacts in the time domain were removed manually. Muscle activity and mechanical artifacts were rejected, while eye movements were kept in the data. The data was then re-referenced to average reference and submitted to independent component analysis (ICA, [30]). An equivalent dipole model for all independent components (IC) was computed using the Boundary Element Model (BEM) based on the MNI brain (Montreal Neurological Institute, MNI, Montreal, QC, Canada) as implemented by DIPFIT routines [31]. ICs reflecting eye movement and one frontal IC reflecting mainly 50 Hz noise were removed based on their scalp maps, component activity, and power spectrum. On average, 5 ICs per participant (*Min =* 2 ICs, *Max* = 11 ICs) were excluded before the data was back projected to the sensor space.

For the analysis of ERPs, the original sensor data was preprocessed using the identical processing steps as described above excluding the data cleaning in the time domain. The respective weights and spheres matrices from the ICA solution were copied to this data set. Afterwards, epochs were extracted from −500ms to 1500ms after stimulus onset of the landmark pictures in the landmark recognition test. The epochs were baseline corrected using the pre-stimulus period from −200ms until stimulus onset. Only trials with correct response were accepted. Epochs of one landmark type (relevant, irrelevant, or novel) were aggregated to one dataset resulting in three datasets per subject. Subsequently, an automatic epoch rejection with a threshold of 100 µV for extremely large fluctuations and a rejection of improbable activity (probability threshold 5 standard deviations and a maximum rejection of 5% of the trials per iteration) was computed. The remaining 126 datasets (3 landmark types x 42 subjects) included on average 40 trials (*SD* = 16 trials, *Min* = 3 trials, *Max* = 67 trials) that were used to compute the averaged event-related potential. For the late positive complex (LPC), an average voltage for a time window from 300 to 700ms was computed. Parietal electrodes (P3, Pz, P4) were selected as the LPC was shown to be most pronounced over posterior regions [16, 17].

### 2.5 Statistical Analyses

Statistical analyses were performed using the statistic software SPSS (International Business Machines Corporation (IBM) Analytics, Armonk, USA). For the analysis of mental load, a 2×3 factorial mixed measures Analysis of Variance (ANOVA) was calculated with the factor navigation session (First and Second Navigation Session) as repeated measure and the factor navigation instruction condition (standard, contrast modified, personal-reference modified) as between subject measure. An additional 3×3 factorial mixed-measure ANOVA was computed focusing on the landmark recognition task using the between-subject factor navigation instruction condition and the within-subject factor landmark type (relevant, irrelevant, and novel). For the analyses of ERP amplitudes, the factor electrode was added to result in a 3×3×3 factorial mixed-measures ANOVA with the between subject factor navigation instruction condition, the repeated measures landmark type and electrode position, including the left, central, and right-hemispheric parietal electrodes (P3, Pz, and P4). Post-hoc tests were adjusted for multiple comparisons with Fisher’s Least Significant Difference (LSD) test. In case of violation of sphericicity, Greenhouse-Geisser corrected values and degrees of freedom are reported. As indicator for the effect size, the partial eta squared was calculated.

## 3 Results

### 3.1 Mental Load

The 2×3 factorial mixed-measure ANOVA revealed no effect of the factor navigation instruction condition 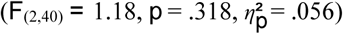 on subjectively experienced mental load (NASA-TLX subscale) of participants. A significant main effect was observed for the navigation session 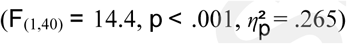 indicating that the overall mental load during the navigation drive without navigation assistance was higher (*M =* 61.6, *SE =* 3.01) than during the navigation-guided drive (*M =* 47.2, *SE =* 3.35). There was no interaction effect of the two factors 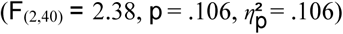.

Driving performance parameters (mean distance to ideal line, mean velocity, number of pedal changes, and number of steering wheel changes) were investigated as objective indicators for mental load during driving. Route segments during wrong turns were excluded from calculations. The 2×3 factorial mixed-measure ANOVAs for the different dependent variables revealed similar statistical effects and thus only the results for the mean distance to the ideal line are reported here. There was no impact of navigation instruction conditions on the mean distance to the ideal line 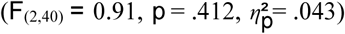 and no interaction effect of navigation instruction condition and navigation session 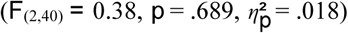 was revealed. There was a significant main effect for navigation session 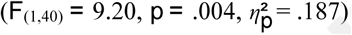 caused by a higher mean distance to ideal line during the second driving session without navigation assistance (*M =* 1.39m, *SE =* 0.02m) as compared to the first driving session (*M =* 1.33m, *SE =* 0.01m).

### 3.2 Performance

In the landmark recognition task, the rates of correct responses for each landmark type were taken as indicator for spatial knowledge acquisition during the assisted navigation session. The percentage of correct responses to relevant landmarks was defined as turning the steering wheel irrespective of the direction. The 3×3 mixed-measures ANOVA revealed a significant main effect of navigation instruction condition 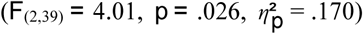 and a significant main effect for landmark type 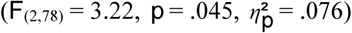.The interaction between navigation instruction condition and landmark type did not reach significance 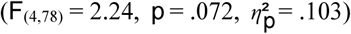. Post hoc comparison revealed significantly lower recognition rates for participants in the standard instruction condition (*M =* 51.7%, *SE =* 2.3%) as compared to both modified navigation instruction conditions (contrast modified: *p =*.034, *M* = 58.9%, *SE =* 2.3%; personal-reference modified: *p =*.012, *M =* 60.7%, *SE =* 2.4%). Post hoc comparisons for the different landmark types revealed higher recognition rates for irrelevant landmarks (*p =*.006, *M =* 64.1%, *SE =* 3.2%) as compared to relevant landmarks (*M =* 50.6%, *SE =* 3.2%). This effect was driven by the pronounced differences between irrelevant and relevant landmarks in the standard navigation instruction condition (see Figure 2). No other differences were significant.

**Figure 2:**
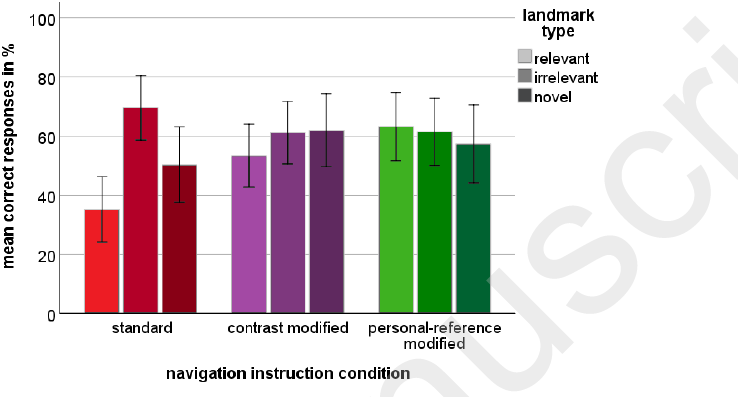
Mean correct responses in % as a function of landmark type and navigation instruction condition. Error bars display +/-2 standard error.

### 3.3 Event-Related Brain Activity

An overview of the event-related potentials with onset of landmark presentation is shown in figure 3 for nine representative channels of the 64 channel montage. While the differences between the navigation instruction conditions became noticeable over posterior recording sites, no such differences were present for different landmarks types.

**Figure 3:**
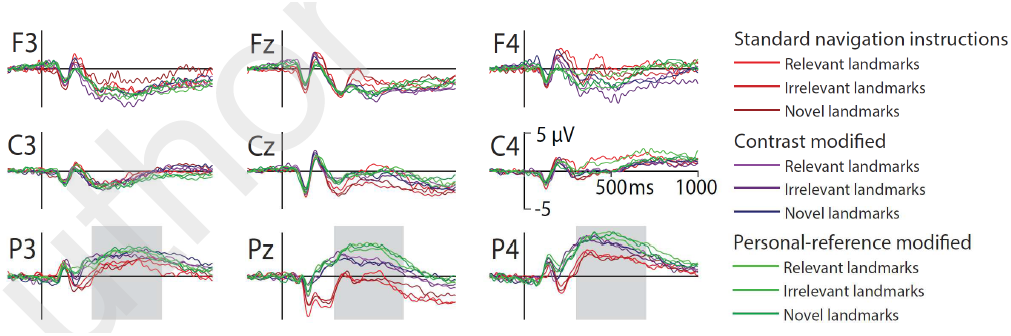
Event-related potentials displayed at frontal (F), central (C) and parietal (P) electrodes over the left hemisphere (3), the midline (z), and the right hemisphere (4). Data is displayed from −200ms until 1000ms of stimulus onset. Navigation instruction conditions and landmark types are color coded with standard navigation instructions in red colors, contrast modified navigation instructions in blue colors, and personal reference navigation instructions in green colors. Landmark types are coded by color intensity. Voltage is displayed with positivity upwards. The grey bar at parietal leads indicates the time window used for statistical analysis of the late positive complex.

The results of the mixed measures ANOVA revealed a significant main effect for the navigation instruction condition 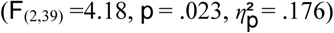. The LPC for standard navigation instructions demonstrated significantly lower amplitudes (*M =* 1.02µV, *SE* = 0.57µV) than the LPC in both modified navigation instruction conditions (contrast condition: *p =*.041, *M =* 2.69µV, *SE* = 0.55µV, personal-reference condition: *p =*.009, *M =* 3.27µV, *SE* = 0.59µV). The main effect of electrode position 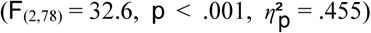 revealed higher LPC amplitudes over the right hemisphere (*M* = 3.37µV, *SE* = 0.33µV) as compared to the midline (*p* <.001, *M =* 1.40µV, *SE =* 0.39µV) and the left parietal leads (*p* <.001, *M =* 2.21µV, *SE* = 0.39µV). These main effects were qualified by a significant interaction effect between navigation instruction condition and electrode position 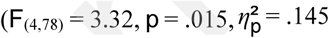 see figure 4). This interaction effect was mainly due to lowest LPC amplitudes but strongest differences between instruction conditions at Pz while the LPC-amplitudes were more pronounced and condition differences less pronounced at the left-parietal lead. At Pz, both modified navigation instruction conditions were associated with significantly increased LPC amplitudes compared to the control group (standard: *M =* −0.50µV, *SE* = 0.68µV; contrast modified: *p* =.020, *M =* 1.78µV, *SE* = 0.65µV; personal-reference modified: *p* =.001, *M =* 2.94µV, *SE* = 0.70µV). For the right-parietal electrode only the difference between personal-reference modification (*M =* 4.23µV, *SE* = 0.56µV) and standard instructions was significant (*p* =.017, *M =* 2.16µV, *SE* = 0.57µV. No effect including the factor landmark type reached significance (*p* >.089).

**Figure 4:**
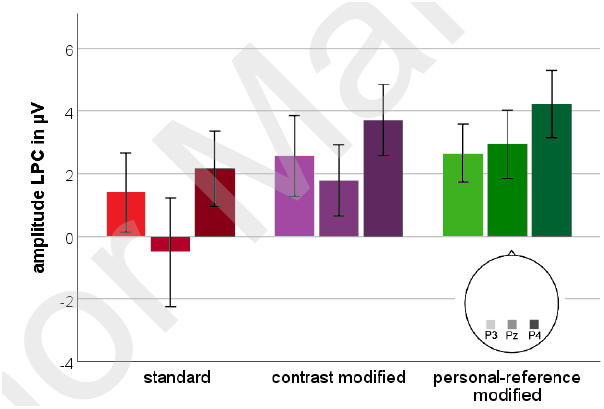
Amplitude of the late positive complex in the time window 300ms to 700ms after stimulus onset for each navigation instruction condition and electrode position (P3, Pz, P4). Error bars display +/-2 standard error.

## 4 Discussion

This study investigated whether modified navigation instructions improve long-term spatial memory for an unknown environment. Participants navigated through a virtual city with the aid of standard or modified navigation instructions. The results demonstrated that, even after being exposed to a new environment only once and with a break of three weeks, landmark recognition performance was significantly improved for participants receiving modified navigation instructions. In addition, electrophysiological data revealed a significant difference between the navigation instruction conditions without differentiating between the landmark types.

### 4.1 Mental Load

To control that the modified navigation instructions did not increase the mental load of the user compared to a standard navigation assistance system, subjective (NASA-TLX) and objective (driving parameters) measurements were investigated. The analyses revealed no differences between the navigation instruction conditions but increased mental load during the second navigation drive that provided no navigation assistance. These results replicate the results of previous studies demonstrating decreased mental load during driving and navigating with the use of navigation assistance systems [32, 33]. Importantly, load measures did not differ for navigation instruction conditions rendering modified navigation instructions a useful option for improving spatial knowledge acquisition in assisted navigation.

### 4.2 Performance

Based on previous results [5], we expected improved landmark recognition performance in the modified navigation instruction conditions. Landmark recognition rates were expected to be best for personal-reference instructions, followed by contrast modified instructions and the standard instructions. Relevant landmarks were expected to be recognized better, because of their navigation decision relevant locations and because only those landmarks were accompanied by an auditory cue specifically mentioning the landmark.

The results of the recognition task showed the expected main effect of navigation instruction condition. A significant higher rate of correctly recognized landmarks was observed for participants in both modified instruction conditions as compared to the control group. These results support the assumption that highlighting an object along the route has a strong impact on spatial learning. Even though the improved landmark recognition effect was most pronounced for the personal-reference instructions, there was no statistical difference between both modified navigation instruction conditions. Thus, the present data cannot further clarify whether the improvement is caused by the mere reference to a landmark or by additional landmark-related information. It can be argued that redundant information already triggers further elaboration [34] and hence the investigated contrast modification might have already led to increased elaboration. These methodological ambiguities, however, do not diminish the result of significantly improved spatial memory for modified navigation instructions.

Against the hypothesis, the main effect of landmark type revealed higher recognition rates for irrelevant landmarks compared to relevant landmarks. This result is similar to previous findings [5] and questions the assumed predominance of relevant landmarks. Possible reasons for this might be found in the setup of the present study. First, driving the simulated car was a multitasking situation with steering as the main task and navigating plus abiding traffic rules as secondary tasks. Turning at decision points might thus require the focus of attention to be directed towards the main task leading to a suppression of processing environmental features. Especially, for the control group this attentional shift was possibly strengthened by the auditory navigation instructions pointing to the “next intersection” instead of an environmental object. Second, the salience and superposition of the projected directional arrow as visual navigation instruction may have had added to this general suppression of processing the surrounding area. Both factors were absent, when passing by irrelevant landmarks and thus, more attention might have been available to process aspects of the environment. This is supported by previous studies reporting reduced recollection of learned words when attention was divided during encoding by a secondary task [35]. Furthermore, passive passengers were reported to recall more landmarks compared to active drivers [36]. Another possible cause for improved recognition rates for irrelevant landmarks, especially in the control group, might have been a response bias. Overall responses revealed significantly more “irrelevant” responses (stepping on the gas pedal) than the chance level for the three response categories would have had predicted^3^. Additionally, incorrect responses were associated significantly more often with pushing the gas pedal than expected. These results supported the presence of a response bias presumably caused by the tendency of participants to respond that a landmark was alongside the route in case of uncertainty. The analyses of recognition performance based on bias-corrected^4^ responses revealed a main effect navigation instruction condition while the main effect of landmark type^5^ could not be replicated.

In conclusion, the attention demanding requirements at navigation-relevant intersections and potential response biases may have negatively impacted the recognition performance for relevant landmarks. Consequently, the higher recognition performance observed in the modified navigation instruction conditions might reflect incidental learning of decision relevant landmarks by neutralizing a suppression effect present in standard navigation instructions.

### 4.3 Brain Activity

The amplitude of the LPC at parietal electrodes is considered an indicator for the recollection process of recognition memory [37]. Recollection is reflected in stronger amplitudes for a higher amount of retrieved information and consequently for “old” versus “new” stimuli. Against the hypothesis, the statistical analyses of the magnitude of the LPC revealed no impact of different landmark types. We expected higher LPC amplitudes for relevant landmarks reflecting improved recognition triggered by additional contrasting or personal relevant information. The results implicate an absence of an “old” (relevant and irrelevant landmarks) versus “new” (novel landmarks) effect. This might be due to the long delay between learning and recognition leading to processes reflecting familiarity judgments rather than recollection of information. However, the generally pronounced amplitudes of the LPC and relatively good performance data contradict that explanation. In another study [38], an old/new effect was absent for stimuli presented with neutral backgrounds and could only be revealed for stimuli encoded within pleasant and unpleasant backgrounds. Thus, the neutral context of the virtual city might have had a similar impact in the present study.

The significant differences between navigation instruction conditions might be explained by a landmark independent recollection of the acquired spatial information. In this case, the onset of each landmark picture triggered the retrieval of context-related information to successfully solve the discrimination task. Thus, the modulation of the LPC amplitude would represent the general amount of retrieved information about the navigated environment instead of landmark-specific information retrieval. It is reasonable to assume that modified navigation instructions altered information processing in a more general rather than landmark-specific fashion as the landmarks were always embedded in the environmental context and not presented in isolation. This effect might have led to higher confidence in participants’ decisions [39]. Thus, larger LPC amplitudes observed in the modified navigation instruction conditions might represent an increased amount of retrieved information covarying with higher confidence due to a generally increased environmental knowledge. A right hemispheric specialization was found in several studies investigating human spatial navigation [40, 41] as well as studies on attention that demonstrated a right-hemispheric asymmetry of activity in the parietal cortex (e.g. [17, 42, 43]). Furthermore, Tucker et al. [44], who examined spatial memory in terms of spatial-attention shift on the visual field and intentional memory of places of stimuli, described this activity of visual search as a negative attenuation of the LPC in the time frame from 400-500ms of the ERP. This negativity was stronger at the contralateral hemisphere for an attentional shift towards one side of the visual field plus a superior spatial activity of the right hemisphere leading to lower LPC amplitudes through superposition [44].

In conclusion, pronounced differences in LPC amplitudes for different navigation instructions but not different landmark types might be explained by less effort required in visual search for relevant environmental features in the modified navigation conditions. This would support the assumption that the amount of recollected information and the confidence of the decision is higher and thus the spatial learning is enhanced in modified navigation instructions.

## 5 Outlook

This study provides a better understanding of spatial learning induced by modified navigation instructions and its underlying brain activity. The advantage of pointing out landmarks alongside the route was replicated providing a scientific basis for integrating modified navigation instructions in future navigation assistance systems to support spatial learning. The influence of emphasizing irrelevant landmarks and providing additional landmark related information should be investigated. Future studies will have to address the concept of the contrast modification including meaningful instead of redundant additional information. Finally, because GPS-based assistance systems are used repeatedly also in well-known environments, it is important to examine the effects of long-term spatial learning based on the repetitive use of modified navigation instructions.

Further studies will provide new insights into the basis of spatial memory and its cortical correlates connected to the use of navigation assistance systems. If such systems succeed to support the neural foundation of spatial cognition and more general memory processes, the implementation of the results in commercially available systems might help postponing dementia-based cognitive declines.

## Acknowledgements

This work was supported by a stipend from the Stiftung der Deutschen Wirtschaft to AW. We would like to thank Matthias Rötting at TU Berlin for providing the car simulator facilities and Sabine Grieger for helping to conduct the experiment.

## Footnotes

1 The test involves sending electrical impulses (n=3600, 50ms average recurrence) from a low-latency interface (the parallel port, controlled via MATLAB) of a PC directly to our ActiCap EEG electrodes. Impulse amplitude is limited to ~ 150 mV via a voltage divider (ratio: 1:20). “At the same time” this PC sends “LSL markers” (string formatted irregularly sampled time series data) over the network, the “ground truth” in this setup. Another PC runs the BrainVisionRecorder (gathering data from the USB adapters), the BrainVision LSL application (“converting” these data samples into an LSL stream), and LSL’s LabRecorder recording the “EEG data” (electrical impulses) and the markers sent from the first PC. The age of sample then is given by the time stamp difference of the markers and the corresponding impulse flanks in the BrainVision data.

2 https://www.optoma.de/projectorproduct/gt1080e (last access: 23.02.2018)

3 The “irrelevant” responses (i.e., stepping on gas pedal) were significantly more often than one third of the trials (*M =* 38.7%, *SE* = 1.8%, *t*_(41)_ = 3.13, *p* =.003). The “relevant” responses (i.e., turning the steering wheel) were observed significantly less than in one third of the trials (*M =* 27.1%, *SE* = 2.1%, *t*_(41)_ = −2.85, *p* =.007). The rate of “unknown” decisions (i.e., stepping on the brake pedal) demonstrated the expected response distribution (*M =* 34.3%, *SE* = 2.5%, *p* =.614).

4 Bias correction was computed via multiplication of the percentage correct with a factor 33.3% divided by mean percentage reaction kind over all conditions respectively for each landmark type.

5 Main effect landmark type: 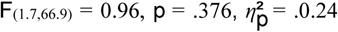 Main effect navigation instruction condition: 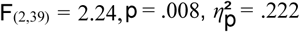, with post hoc comparison showing a significant difference between standard (*M* = 51.5%, *SE* = 2.5%) and contrast condition (*p =*.013, *M* = 60,6%, *SE* = 2.4%) and personal-reference condition (*p =*.004, *M* = 62,7%, *SE* = 2.6%). The interaction effect of navigation instruction condition and landmark type is still not significant 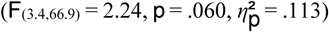.

